# Characterization of heterogeneity in nanodisc samples using Feret signatures

**DOI:** 10.1101/2022.07.28.501900

**Authors:** Fernando Vilela, Armel Bezault, Borja Rodriguez de Francisco, Cécile Sauvanet, Xiao-Ping Xu, Mark F. Swift, Yong Yao, Francesca M. Marrasi, Dorit Hanein, Niels Volkmann

## Abstract

Nanodiscs have become a popular tool in structure determination of membrane proteins using cryogenic electron microscopy and single particle analysis. However, the structure determination of small membrane proteins remains challenging. When the embedded protein is in the same size range as the nanodisc, the nanodisc can significantly contribute to the alignment and classification during the structure determination process. In those cases, it is crucial to minimize the heterogeneity in the nanodisc preparations to assure maximum accuracy in the classification and alignment steps of single particle analysis. Here, we introduce a new *in-silico* method for the characterization of nanodisc samples that is based on analyzing the Feret diameter distribution of their particle projection as imaged in the electron microscope. We validated the method with comprehensive simulation studies and show that Feret signatures can detect subtle differences in nanodisc morphologies and composition that might otherwise go unnoticed. We used the method to identify a specific biochemical nanodisc preparation with low size variations, allowing us to obtain a structure of the 23-kDa single-span membrane protein Bcl-xL while embedded in a nanodisc. Feret signature analysis can steer experimental data collection strategies, allowing more efficient use of high-end data collection hardware, as well as image analysis investments in studies where nanodiscs significantly contribute to the total volume of the full molecular species.

**GRAPHICAL ABSTRACT:** 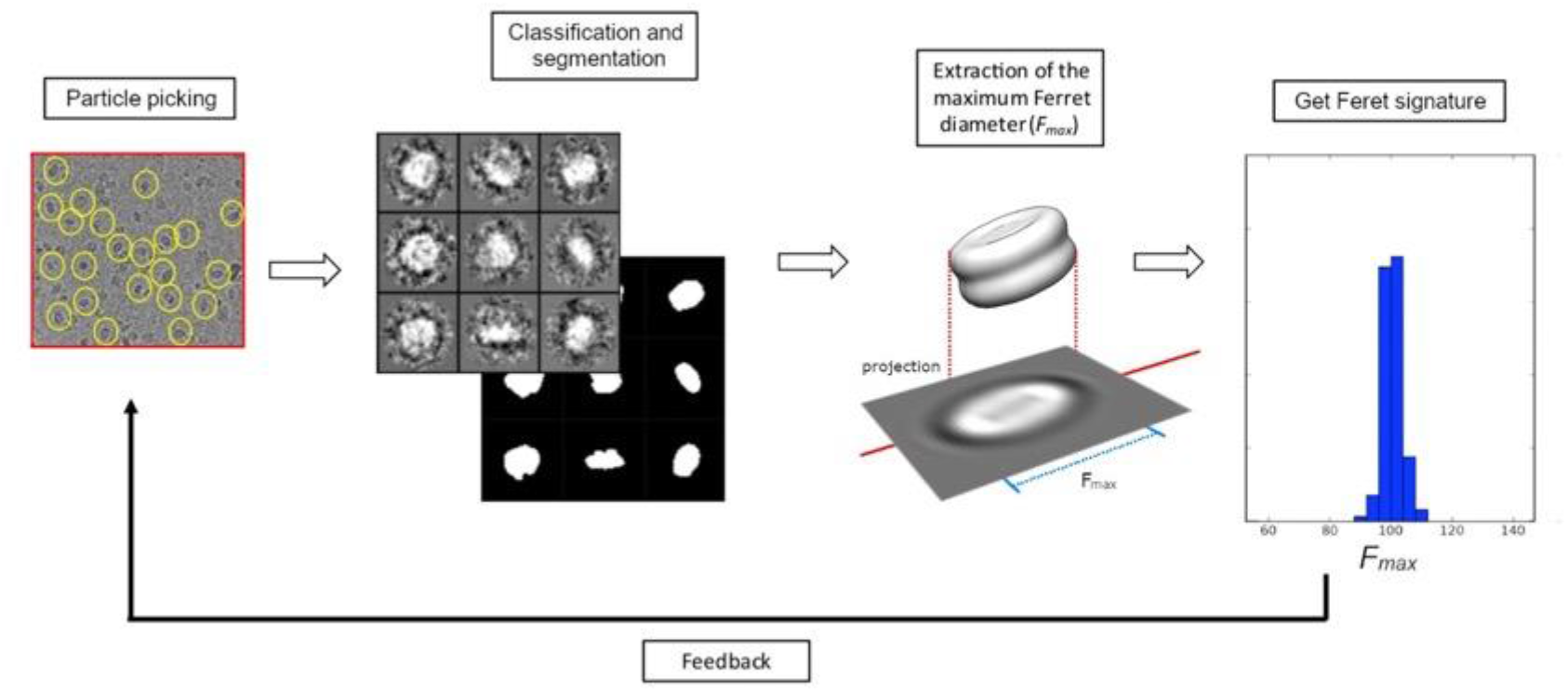

**HIGHLIGHTS:**

- New methodology to characterize nanodiscs based on Feret signatures
- Feret signatures distinguish nanodisc morphologies and compositions
- Analysis is highly sensitive to sample quality
- Method selected condition to solve structure of small membrane protein Bcl-xL

## INTRODUCTION

Membrane proteins play key roles in signaling, transport, and metabolism. Structural studies of membrane proteins are typically challenging, owing to the need to include a proper lipidic environment that reproduces their physiological state as closely as possible. In recent years, the development of nanodiscs and related multi-molecular systems, which provide membrane proteins with a controlled lipidic environment, have led to major advances. These multi-molecular systems are supported by belts that contain and maintain a stable lipid bilayer and render them soluble and stable in aqueous solutions. Notably, membrane scaffold protein (MSP) nanodiscs (McLean et al., 2018), steryne-maleic acid lipidic particles (SMALPs) forming co-polymer nanodiscs (Dörr et al., 2016) and covalently circularized nanodiscs (cND) (Nasr et al., 2016) serve as controlled lipidic environment for membrane proteins. While this study focuses on MSP nanodiscs, the general principles of the protocols described are applicable to any disc-like system including, among others, SMALPS and cNDs. Because detergents often modify the native conformation and function of membrane proteins, an important advantage of nanodiscs is that they are detergent free.

MSPs are amphipathic molecules derived from the apolipoprotein-1 sequence that. Two MSP molecules self-assemble as a belt, wrapping around lipids and membrane proteins to form discoidal nanodiscs. A wide range of experiments and studies have been realized using MSP nanodiscs (Sligar and Denisov, 2020; Denisov and Sligar, 2017). In particular, the combination of nanodiscs and Single Particle Analysis by cryogenic electron microscopy (cryo-EM) allows near atomic information to be achieved for membrane proteins closer to their native physiological state (Efremov et al., 2017; Cheng, 2018). Single spanning (or bitopic) membrane proteins represent more than half of the membrane protein in analyzed genomes but only a small fraction of the solved membrane protein structures constitute single spanning membrane proteins (Allen et al., 2019). The conformation of single spanning membrane proteins can be strongly influenced by the membrane environment (Xu et al., 2016) so nanodiscs are an attractive possibility for structural analysis of these proteins. However, these proteins tend to have low molecular weight (< 100kDa) and are thus in the same size range as nanodiscs. In contrast to large multi spanning membrane proteins, in the structural studies of low molecular-weight membrane proteins the nanodisc itself will significantly contribute to the classification, alignment and reconstruction process. Thus, it is essential to minimize the heterogeneity in nanodisc preparations when studying small single - spanning membrane proteins.

Here, we describe a robust and accurate protocol for systematically identifying and classifying heterogeneity in nanodisc samples based on a new sample characterization paradigm we coined Feret signature analysis. This analysis can steer experimental data collection strategies, allowing more efficient use of high-end data collection hardware, as well as image analysis investments. The protocol aims to overcome bottlenecks to achieving high-resolution structures of single spanning and low molecular-weight membrane proteins in nanodiscs where nanodiscs significantly contribute to the total volume of the full molecular species (nanodiscs with incorporated membrane proteins).

We demonstrate the advantages of this protocol for the structural analysis of a 23-kDa single-spanning membrane protein incorporated in nanodiscs: the human Bcl-2 family protein Bcl-xL. Bcl-xL is a major suppressor of apoptosis, a fundamental homeostatic process of programmed cell death that is conserved across a wide range of organisms and plays key roles in major human diseases. The protein is expressed in a variety of tissues and cell types and overexpressed in many tumors where it acts to promote tumor cell survival, tumor formation and tumor resistance to chemotherapy. Like most other members of the Bcl-2 protein family, Bcl-xL functions at intracellular membranes and its C-terminal membrane-anchoring segment is critical for function. Nevertheless, structural studies have advanced primarily by deleting its membrane anchor and adding detergents to enhance solubility. As a result, the molecular mechanisms controlling the intracellular localization and cytoprotective activity of Bcl-xL are not completely understood. Recently, we used NMR spectroscopy to describe the conformations of Bcl-xL with intact C-terminus (Yao et al., 2015) and full-length Bcl-xL (Ryzhov et al., 2020) in detergent-free lipid bilayer nanodiscs. In this state, the protein is anchored in the nanodisc membrane by its helical tail while the N-terminal head retains the canonical solvent-exposed state that is competent for ligand binding. Notably, membrane-associated Bcl-xL has enhanced BH3 binding affinity compared to its tail-truncated, soluble counterpart, indicating that the membrane provides an additional level of regulation and underscoring the importance of understanding protein structure and function in the membrane environment.

We validated the protocol with simulated cryo-EM samples containing bare nanodiscs (MSP nanodiscs in the absence of membrane proteins) and demonstrated its use with experimental bare nanodisc samples. In addition, we identified nanodisc preparations that allowed successful structure determination of Bcl-xL with intact C-terminus embedded in nanodiscs, using our protocol.

## MATERIAL AND METHODS

### Simulated Data

Nanodiscs were modeled on a three-dimensional 400 × 400 × 400 grid with 0.5 Å spacing. Voxels within the shape were set to one and voxels outside the shape were set to zero. This operation was followed by Gaussian smoothing with a box width of 5 voxels. Random rotations were applied by picking a rotation vector from an equidistant distribution of points on a sphere (Saff and Kuijlaars, 1997), applying the corresponding rotation vectors and then adding a random rotation around that vector. Defocus was applied to the volumes using the formulation by Thon (Thon, 1971) using the weak phase object and projection approximations. Projections were generated by summing all Z-sections and subsequent binning to 4 Å pixel size. A mix of correlated Gaussian and Laplacian random noise was used to approximate noise of experimental nanodisc data sets. Mixing and correlation parameters were adjusted to match the distributions and texture of the experimental noise. When adding to the simulated projection data, the scale factor was adjusted to generate images with signal-to-noise ratio similar to the experimental data.

### Reconstitution and purification of bare nanodiscs and Bcl-xL-nanodisc assemblies

Protein purification and nanodisc preparation were performed as described (Ritchie et al., 2009; Hagn et al., 2013; Yao et al., 2015; Yao and Marassi, 2019). Briefly, the gene (Genbank NM_001191) encoding human Bcl-xL, but lacking residues 44-84 in the flexible loop, was cloned and expressed in *E. coli* BL21 cells and purified by chromatography. The MSP variants MSP1D1Δh5 and MSP1D1, were expressed in *E. coli* and purified following the established protocols. All lipids (Avanti Polar Lipids) were taken from chloroform solution and dried under vacuum overnight. Bcl-xL was incorporated into MSP1D1Δh5 and MSP1D1 nanodiscs prepared with a 3:1 molar mixture of dimyristoyl-phosphatidyl-choline (DMPC) and dimyristoyl-phosphatidyl-glycerol (DMPG) phospholipids. Three solutions were prepared and then combined: (i) pure lyophilized Bcl-xL was dissolved in nanodisc buffer (20 mM Tris-Cl, pH 7.5, 2 mM DTT, 1 mM EDTA) containing 100 mM n-decyl-phosphocholine (DePC; Anatrace); (ii) DMPC and DMPG were co-dissolved in 1 mL of nanodisc buffer containing Na-cholate (Anatrace) to obtain a final 1:2 molar ratio of lipid to cholate; and (iii) purified MSP was dissolved in nanodisc buffer. The combined solution was incubated at room temperature for 1 hr and then mixed with BioBeads SM-2 resin (BioRad) overnight to remove the detergents. The BioBeads were removed by centrifugation and the resulting nanodiscs were washed twice with one sample volume of nanodisc buffer. The nanodisc solution was transferred to EM buffer (25mM Sodium phosphate, pH 6.5, 50mM NaCl, 2mM DTT and 1mM EDTA), concentrated using a 10 kD cutoff Vivaspin concentrator (Viva Products) and stored at 4°C. The MSP:lipid ratio was adjusted to 1:50 in the case of MSP1D1Δh5 and 1:60 in the case of MSP1D1. The nanodisc homogeneity was analyzed by size exclusion chromatography. Bare MSP1D1Δh5 and MSP1D1 nanodiscs with DMPC and DMPG were prepared following the same protocol without Bcl-xL.

MSP1D1 nanodiscs were also reconstituted with palmitoyl-oleoyl-phosphatidyl-choline (POPC). POPC was solubilized in nanodisc buffer incremented with 70 mM Na.Cholate. MSP1D1 and POPC were mixed at a molar ratio of 1:33 (MSP1D1:POPC). The mixture was incubated 1h at 5°C, with 12 rpm agitation. Then, BioBeads SM-2 resins were added in excess, aiming at the complete removal of Na.Cholate from the mix. The Nanodiscs reconstitution samples were incubated overnight at 5°C and 12 rpm agitation. Then, BioBeads were removed, samples were filtered at 0.2 μm, and nanodiscs were purified by size-exclusion chromatography, using a Superdex 200 increase 10/300 GL 24 mL, where approximately 500 μL of each nanodisc sample were injected. Gel filtration experiments were performed at 0.35 mL/min, 5°C, and relevant elution fractions were collected. The purity of nanodiscs was confirmed by SDS-PAGE 15% and fractions were selected to be concentrated by Vivaspin 500 Centrifugal Concentrators (Sartorius, ref. VSO0112) with centrifugation cycles of 5 min, 10.000 rpm, at 5°C. Finally, UV-Vis spectra were acquired and nanodisc concentration was calculated by absorbance at 280 nm.

### Dynamic Light Scattering

Dynamic light scattering (DLS) was performed to characterize bare nanodisc MSP1D1:POPC samples using a Wyatt Technology DynaPro Plate Reader III instrument at the scattering angle of 169°, with a laser wavelength of 830 nm. Nanodiscs samples were prepared at 0.3 mg/mL and centrifuged before analysis. A volume of 20 µL of sample was loaded in a 384-well microplate (Corning ref 3540, New-York, USA), with 20 acquisitions of 10 s each at 20°C, monitored with the DYNAMICS version V7.10.0.21 software (Wyatt, Santa Barbara, CA, USA), 3 repetitions per measurement. Experimental design and analysis of the experiments were realized using the same software.

### Sample vitrification

4 μL of Bcl-XL-nanodisc assemblies and bare MSP1D1Δh5 nanodiscs were applied on freshly glow-discharged Quantifoil copper 400 mesh 2/1 holey carbon film cryo-EM grids, manually blotted, and plunged into liquified ethane at LN_2_ temperature using an in-house designed Cryo Plunger at a humidity between 90 and 100%. MSP1D1 nanodiscs were applied on freshly glow-discharged Quantifoil copper 400 mesh 1.2/1.3 holey carbon film and were vitrified using a VitroBot IV automatic plunger (ThermoFisher) at 4°C and 100% humidity, with blot time 3 s, waiting time 30 s, and blot force 2. Long-term storage of grids was done in liquid nitrogen.

### Cryo EM imaging

Sample grid quality was assessed before data collection using a Tecnai Sprit T12 microscope (ThermoFisher) or a Glacios (ThermoFisher). Cryo-EM data sets were collected on a Titan Krios electron microscope (ThermoFisher) operated at 300 kV acceleration voltage with a Falcon II direct electron detector (ThermoFisher) or were acquired on a Glacios electron microscope (ThermoFisher) operated at 200 kV acceleration voltage, with a Falcon 4i direct electron detector (ThermoFisher) in EER mode. Automated data collection was carried out with EPU software (ThermoFisher) with calibrated pixel size between 1 and 3 Å at defocus ranges between 1.7 and 4.2 μm and with a total dose between 50 and 60 e^-^/Å^2^.

### Data processing

The frames were aligned, dose-weighted, and summed using MotionCor2 (Zheng et al., 2017). The CTF of the individual images was estimated with CTFFIND4 (Rohou and Grigorieff, 2015). Particle picking for the MSP1D1Δh5 data sets was done manually with EMAN2 (Tang et al., 2006). Particle picking for the MSP1D1 data set was done automatically with RELION (Scheres, 2015). Pixel sizes were reduced for the extracted particles by binning to a value between 4 and 5 Å and boxes were cropped tightly for Feret analysis. Particle picking for the Bcl-xL-nanodisc assembly data sets was done manually with RELION. Reconstructions of the Bcl-xL-nanodisc assemblies were generated following the RELION workflow. The final resolution reported by RELION for the Bcl-xL-MSP1D1Δh5 assembly was 15Å. The value for the nanodisc thickness *t* (57 Å) used for calculation of *d*_*nd*_ was taken from small angle X-ray scattering analyses (Denisov et al., 2004). P-values were calculated with a two-sided t-test.

### Characterization of nanodisc distributions

*F*_*max*_ signatures were extracted following a multi-step procedure composed of (1) reference-free classification and averaging, (2) denoising, (3) binarization, (4) segmentation, and (5) extraction of the maximum Ferret diameter. For noise-free simulated date, the first two and the fourth steps were omitted. Because of the simple disc shape of nanodiscs without discernible internal features, it is not necessary to use highly sophisticated and computationally expensive approaches such as those based on maximum likelihood for the reference-free classification step. We found that an approach based on rotationally and translationally invariant double autocorrelation functions (Schatz and van Heel, 1990) performs just as well in the context of nanodisc classification with about an order of magnitude improvement in classification speed.

In addition to the classification/averaging step, we introduced a noise reduction step based on non-local means (Buades et al., 2005) to further enhance the signal-to-noise ratio. Binarization between foreground and background is based on maximizing the sum of their Renyi entropies of order two (𝓡), subject to modifying the threshold *T* (Yen et al., 1995; Sezgin and Sankur, 2004):

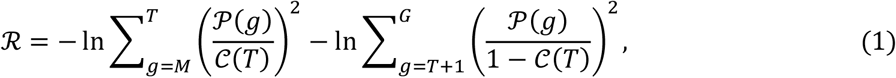

with *g* denoting a grey value at a particular pixel. The first sum is running over all pixels with gray values between the minimum *M* up to the threshold *T*, and the second sum is running over all pixels with gray values larger than *T* up to the maximum gray value *G*. 𝒫 denotes probability estimated from the gray-level histogram. 𝒞(*T*) denotes the cumulative probability of *T*. Conceptually, this algorithm detangles two different signal sources, one coming from the nanodisc density and the other one from background noise.

In rare cases, where the signal-to-noise ratio is still relatively low after classification, averaging, and denoising, the binarization is imperfect and generates additional density blobs outside the nanodisc density. To address this issue, all self-contained blobs are labeled using the watershed transform (Volkmann, 2002) and only the most central one is kept. The entire procedure has been incorporated into *pyCoAn* (github.com/pyCoAn/distro), an extended python version of the CoAn package (Volkmann and Hanein, 1999).

## RESULTS AND DISCUSSION

Availability of a robust technology for determination of size distributions of nanodisc preparations would provide the means to quickly screen the readiness of particular nanodisc batches for pursuing structural studies and, as importantly, provide feedback for refining sample preparation protocols for improving sample homogeneity. This is important not only for cryo-EM but also for solution NMR. Methods based on size exclusion chromatography, dynamic light scattering, or other biophysical approaches used routinely for nanodisc characterization (Efremov et al., 2017; Hagn et al., 2013) focus on bulk (average) characterization and thus are not sufficiently sensitive for capturing and monitoring the full extent of heterogeneity at high resolution. In addition, these biophysical methods characterize the sample in solution, When the sample is applied to the EM-grid and subsequently vitrified, substantial changes may occur including partial denaturation owing to interactions with the air-water interface or selection of particular conformers or multimers during the blotting process. Therefore, it would be preferable to characterize the nanodisc sample after it was applied to the EM grid and it was vitrified.

Standard particle classification protocols of nanodisc cryo-EM data can be misleading, even when performed in three dimensions. For example, class averages derived from elliptical discs or from a mix of discs with two different diameters can be essentially indistinguishable, especially in the presence of noise and if top views are missing (**Figure 1**). However, owing to the unique shape characteristics of the nanodiscs, their size distribution can be used to infer whether the population is fluctuating randomly around a certain size and how much, whether the sample contains nanodiscs with more than one well-defined diameter, or whether it contains elliptical nanodiscs.

**Figure 1:**
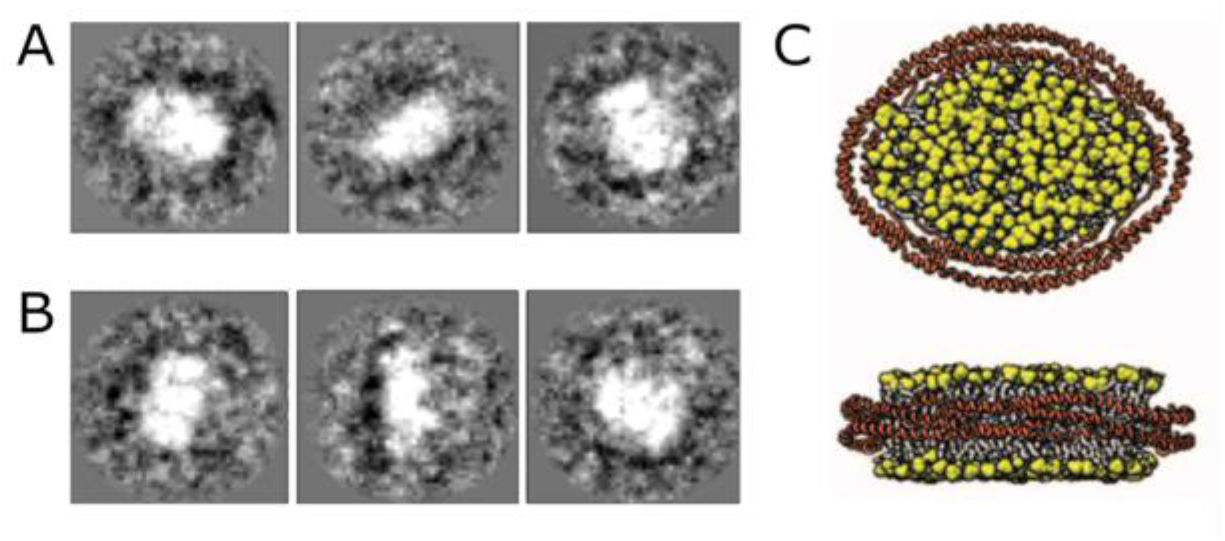
Nanodisc shapes are hard to deduce from noisy projections. (A) Simulated projection data from elliptical nanodisc in the presence of noise and with top views omitted. (B) Simulated projection data from nanodiscs with varying diameter in the presence of noise and with top views omitted. (C) Two orthogonal views of elliptical nanodisc shape used to generate the data in (A).

The inherent problem when relying on purely classification-based methods to deduce nanodisc shapes is rooted in the fact that different objects in different orientations can lead to the same orthogonal projection onto a plane. For example, the orthogonal projection of a tilted circle is an ellipse, which is indistinguishable in shape from the projection of an un-tilted ellipse that matches the projection’s semi-axes. Thus, the occurrence of an elliptical class average by itself will not allow to infer whether the shape of the underlying object is circular or elliptical. This issue is particularly troublesome if some views are systematically missing in a data set, if there are mixtures of different sizes and shapes, or if the signal-to-noise ratio is low, as is common in cryo-EM.

For an infinitely thin disc (i.e. a two-dimensional circle), the diameter can be determined by simply measuring the longest dimension of the disc’s projection image. This corresponds to the maximum Feret diameter *F*_*max*_ of the projection. Feret diameters of an object are defined as the distance between two parallel tangents of the object’s boundaries. In particular, the maximum and minimum Feret diameters, *F*_*max*_ and *F*_*min*_, are often used for the characterization of particle sizes and their distribution in powder samples or polycrystalline solids (Walton, 1948). In the remainder of the manuscript, we refer to the distribution of Feret diameters as ‘Feret signature’ of the sample with ‘*F*_*max*_ signature’ referring to the distribution of the maximum Feret diameter. For an infinitely thin disc, *F*_*max*_ is independent of the disc’s orientation. The reason is that the orthogonal projection of a circle onto a plane is always an ellipse (or a line segment in the degenerate case) with the length of the major axis matching the diameter of the projected circle. For noise-free discs of such type, the *F*_*max*_ signature would be a delta function at the diameter of the disc. A mix of discs with two different diameters would lead to an *F*_*max*_ signature with two delta functions at the two diameters. In contrast, projections of an elliptical disc would lead to a more complex *F*_*max*_ signature that is a function of the major and minor axes *a* and *b*, following *d(θ) = 2 ab / [b* cos*(θ)*^*2*^ *+ a* sin*(θ)*^*2*^]^*1/2*^, with the origin at the center of the ellipse and *θ* being the angle measured from the major axis *a* plus 90 degrees. Thus, *F*_*max*_ signatures of specific samples can be used to infer the shape and composition of the underlying circular/elliptical disc-like objects.

However, nanodiscs have finite thickness and the cryo-EM image formation process introduces various imaging artifacts that need to be considered in an analysis of *F*_*max*_ signatures. In order to test whether *F*_*max*_ signatures are capable of detecting differences between the underlying nanodisc shapes under real-world conditions, we generated simulated data sets derived from simulated nanodiscs. We used a thickness of 48 Å throughout and varied the diameter of the discs between 96 and 136 Å, a typical range for nanodiscs formed with MSP belt proteins (Ritchie et al., 2009). We generated artificial single particle data sets by applying random rotations around the three orthogonal axes to the simulated nanodisc and then projecting the density down the Z axis. This operation was followed by automated thresholding (Yen et al., 1995) and rapid determination of the *F*_*max*_ of the thresholded projection using their moments of inertia. Introduction of the disc thickness *t* and the thresholding/inertia procedures lead to a slight peak broadening of the *F*_*max*_ signature and shifts the mean of *F*_*max*_ (*F*_*mean*_) from the nanodisc diameter *d* to larger values. For top views *F*_*max*_ will still be equal to *d* but for side views *F*_*max*_ is 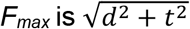. An estimate for the nanodisc diameter *d*_*nd*_ can then be retrieved from *F*_*mean*_ as 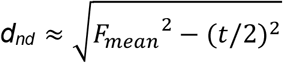.

Because in cryo-EM images the contrast between the buffer and the sample is low, phase contrast is introduced by taking defocused images. Another option is the use of phase plate devices, which enhance contrast while allowing in-focus imaging (Danev and Baumeister, 2016). We run simulations with a fixed nanodisc size of 96 Å and different defocus values, including zero defocus emulating a phase plate data set, and did not see any significant biasing in the *F*_*max*_ signature. Thus, for the rest of the simulations, we used a single defocus of -2.5 μm.

The electrons used for imaging the sample also damage the sample. Thus, the electron dose used for imaging must be kept low. Consequently, the signal-to-noise ratio for the individual images tends to be very low. We introduced random noise into the simulations that closely mimics the appearance of experimental data. The presence of noise critically hampers the accuracy of the thresholding operation and does not allow extraction of an accurate *F*_*max*_ estimate from individual projections. In single particle analysis, classification and averaging are used to improve the signal-to-noise ratio to allow structure determination. Here, we use a similar strategy to get accurate *F*_*max*_ estimates from the noisy data. Clearly, there will be a trade-off between the number of particles contributing to each class and the magnitude of differences between distributions that can be picked up.

For a sample of nanodiscs with well-defined diameter, the generation of class averages and denoising of noisy projections does not significantly alter the *F*_*max*_ signature (**Figure 2**): the *d*_*nd*_ are 95.7 ± 3.7 Å and 95.4 ± 3.3 Å respectively, *p*-value *=* 0.072. Note that both *d*_*nd*_ values are less than 1 Å off the ground-truth value of 96 Å. As a test for more complex scenarios, we checked if we could pick up the difference between simulated noisy data sets of discs with randomly varying diameter between 96 and 136 Å, an elliptical disk with major axes of 96 and 136 Å, and a mixture of two discreet disks with diameters of 96 and 136 Å respectively. Even if we need to increase the class size to 50 particles to combat the noise, we can still pick up the difference (**Figure 3**), showing that the method is useful in a practical setting. Because of the simple shape of discs and the negligible effect of defocusing, it is an advantage to use more strongly defocused images for analyzing the shape distribution because the increased phase contrast will translate to better signal-to-noise ratios. Another useful intermediate step, if data was acquired at higher magnification, is to reduce the pixel size to 4-5 Å, which will also increase contrast through spatial averaging and significantly decreases the time required for the analysis.

**Figure 2:**
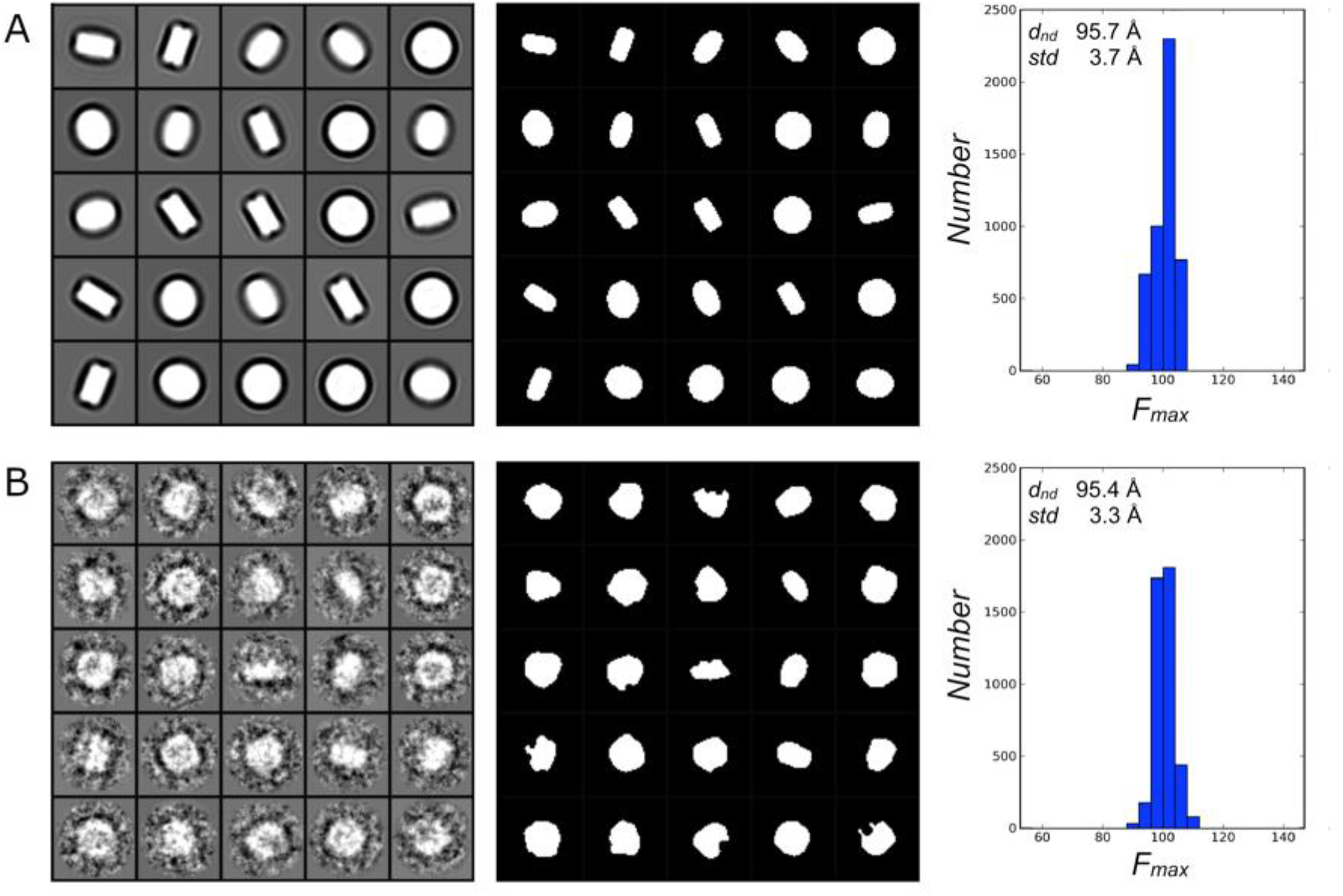
*F*_*max*_ signatures of simulated nanodiscs of 96-Å diameter and 48-Å thickness. (A) *Left panel*: sample projection images of randomly oriented nanodiscs with CTF applied. *Central panel:* binarized projections after segmentation. *Right panel:* distribution of *F*_*max*_ (in Å) extracted from binarized projections. (B) *Left panel*: sample class averages (50 images per class) of projection images of randomly oriented nanodiscs with CTF applied and noise added. *Central panel:* binarized class averages after denoising and segmentation. *Right panel:* distribution of *F*_*max*_ extracted from binarized class averages.

**Figure 3:**
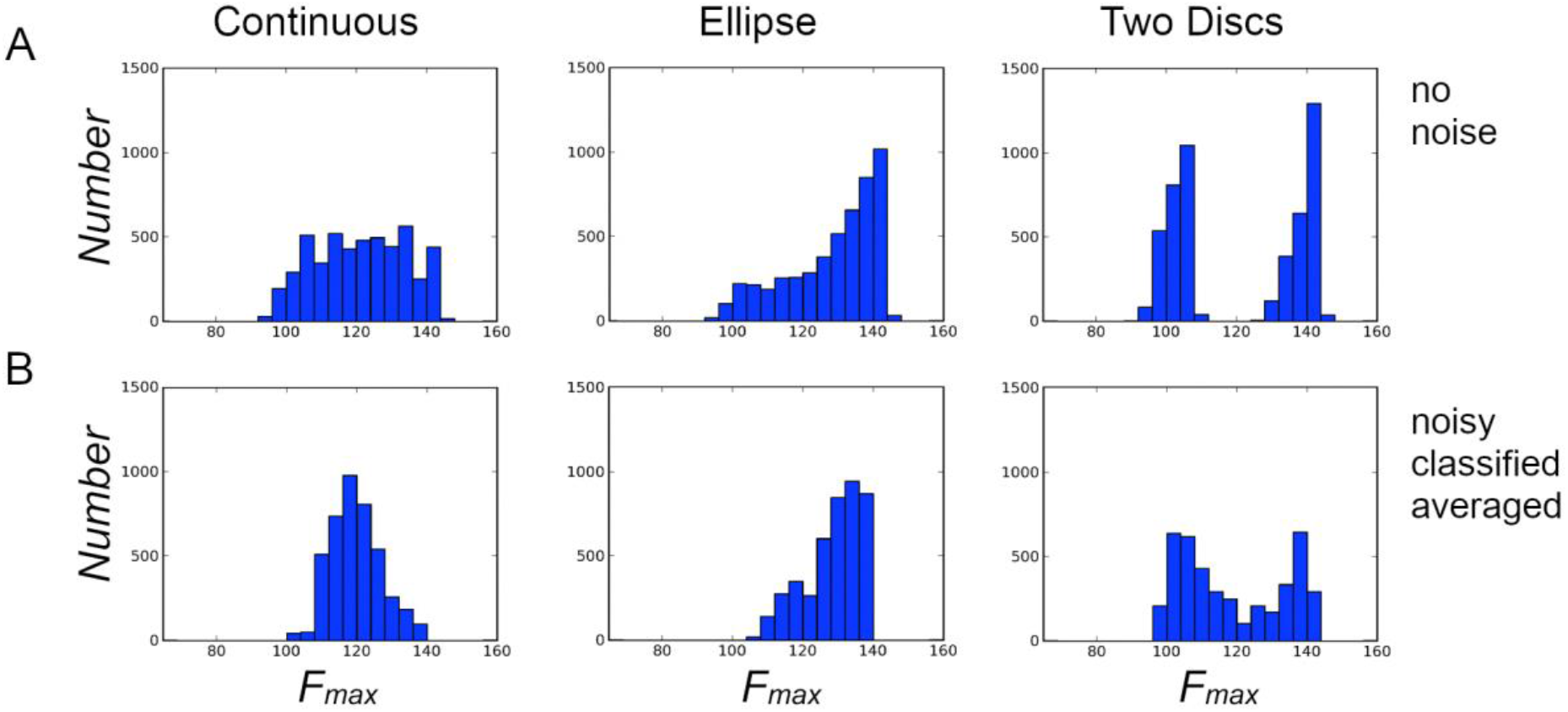
*F*_*max*_ distributions (in Å) of particles randomly picked from different simulated nanodisc shapes and compositions. Thickness of the discs is 48Å for all cases. (A) *F*_*max*_ calculated from individual particle projections in the absence of noise (B) *F*_*max*_ calculated from noisy projection that were classified reference-free into classes with 50 members and then averaged. The left panels (Continuous) were drawn from a nanodiscs with continuously varying diameter between 96Å and 136Å. The center panels (Ellipse) were drawn from elliptical nanodiscs with major axes of 96Å and 136Å. The right panels (Two Discs) were drawn from two distinct nanodiscs of diameters 96Å and 136Å.

Next, we characterized the size distributions of experimental data collected from nanodisc preparations produced with different belt proteins. Namely, we used MSP1D1Δh5 and MSP1D1 (see **Table 1** for nomenclature and size estimates). The MSP1D1 size distribution was first analyzed in solution using dynamic light scattering (DLS). The same sample was then applied to EM grids, vitrified, and analyzed using the Feret signatures (**Figure 4A, B**). The average nanodisc diameter corresponds well (96 Å vs. 97.4 Å) but the DLS-based distribution is much wider than the Feret-based distribution (standard deviation 20.6 Å vs. 11.6 Å), clearly demonstrating the limitations in detectability of subtle differences such as those apparent in Feret distributions of two independent biochemical preparations of MSPD1Δ5 (**Figure 4C, D**). One preparation of the

**Table 1:** Description MSP nanodiscs used in this study (Ritchie et al., 2009). Diameter sizes indicated correspond to stokes hydrodynamic diameters of nanodiscs formed with the corresponding MSPs, determined by size-exclusion chromatography (Denisov et al., 2004) or (marked by asterisk), by dynamic light scattering (Hagn et al., 2013). MW: Molecular weight (monomer).

| Protein | Diameter Size [nm] | MW [Da] | Description |
| --- | --- | --- | --- |
| MSP1 | 9.7 | 24608 | Original MSP1 (deletion 1–43 mutant of Human Apo A-1) |
| MSP1D1 | 9.6 | 24662 | Deletion mutant (residues 1-11) of MSP1 |
| MSP1D1Δ5h | 8.4* | 22106 | Variant of MSP1D1 with deletion of helix 5 |

**Figure 4:**
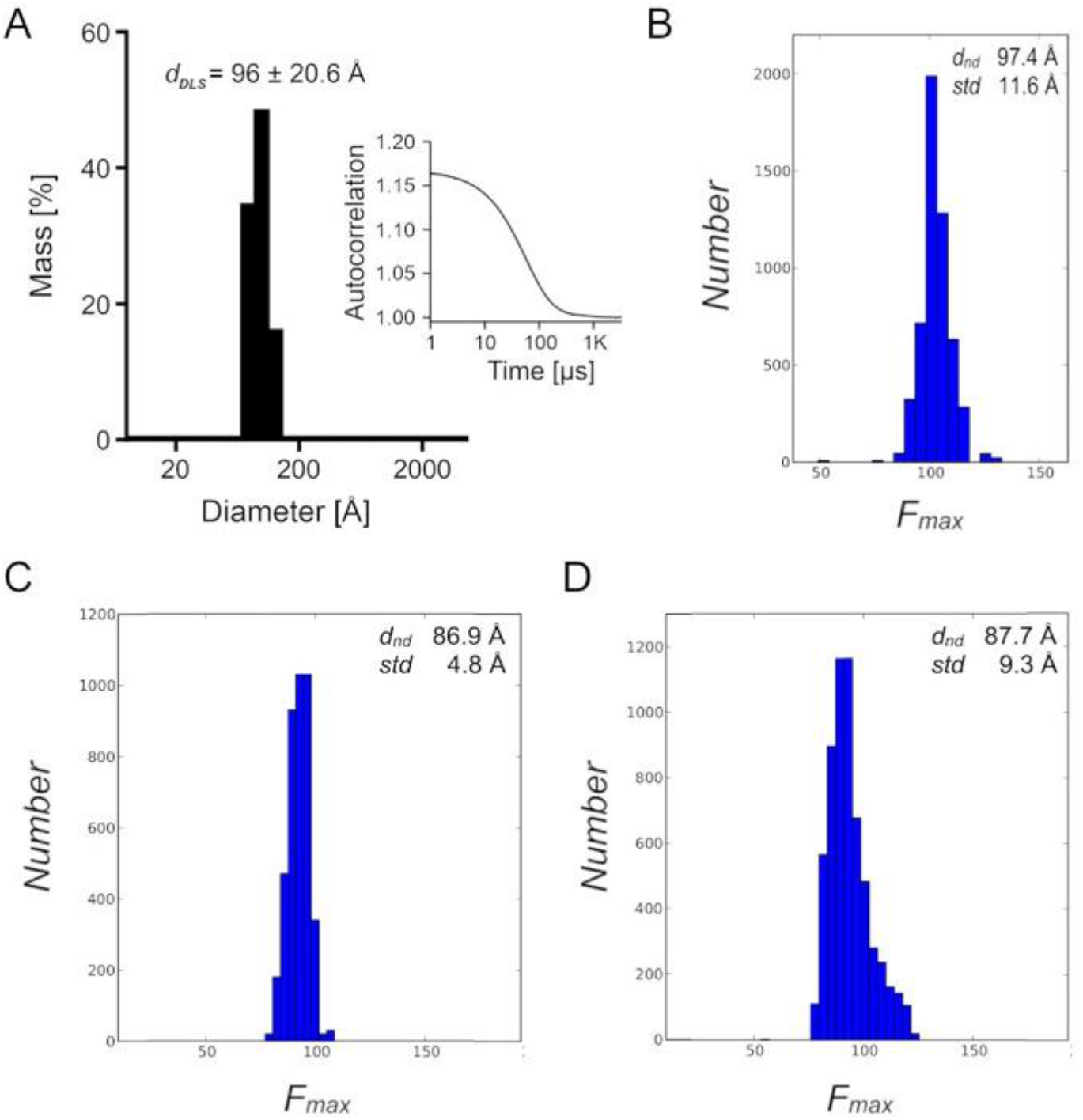
Size distributions of experimental nanodisc samples. (A, B) Analysis of MSP1D1 nanodisc size distribution using dynamic light scattering (DLS) and Feret signature analysis (B). While the mean diameter estimates correspond well, the spread of the Feret distribution is much narrower than that of the DLS distribution. (C, D) *F*_*max*_ distributions of two independent biochemical preparations of MSP1D1Δ5h nanodiscs. The sample in (B) was successfully used to resolve Bcl-xL. MSPD1Δ5 gives a distribution (*d*_*nd*_ = 86.9 ± 4.8 Å) most consistent with a single, well-defined disc diameter (**Figure 4C**) while the other one shows clear signs of peak broadening (*d*_*nd*_ = 87.7 ± 9.3 Å), most consistent in shape with the existence of multiple diameters around the peak value (**Figure 4D**).

We used the MSP1D1Δh5 nanodisc preparation (Hagn et al., 2013) with the narrow size distribution and MSP1D1 nanodiscs as basis for cryo-EM structure determination of anti-apoptotic Bcl-xL. Bcl-xL inserts its C-terminal helix into the membrane while the soluble domain adopts a globular conformation (Yao et al., 2015; Ryzhov et al., 2020) that should be readily visible within Bcl-xL-nanodisc reconstructions (Volkmann et al., 2014). Using the MSP1D1Δh5 nanodisc preparation with the narrow size distribution, we do indeed succeed in determining a structure at an intermediate resolution of 15 Å that clearly shows the Bcl-XL soluble domain on top of the nanodisc (**Figure 5**), confirming the structural information we previously obtained by NMR (Yao et al., 2015; Ryzhov et al., 2020). However, when using MSP1D1 nanodiscs, we failed in deriving a structure that shows the Bcl-xL soluble domain, even at low resolution. The most likely cause is the larger size combined with higher variability of the nanodisc diameter (*d*_*nd*_ = 86.9 ± 4.8 Å for MSP1D1Δh5 versus *d*_*nd*_ = 97.4 ± 11.6 Å for MSP1D1), which disrupts alignment and classification in the reconstruction process.

**Figure 5:**
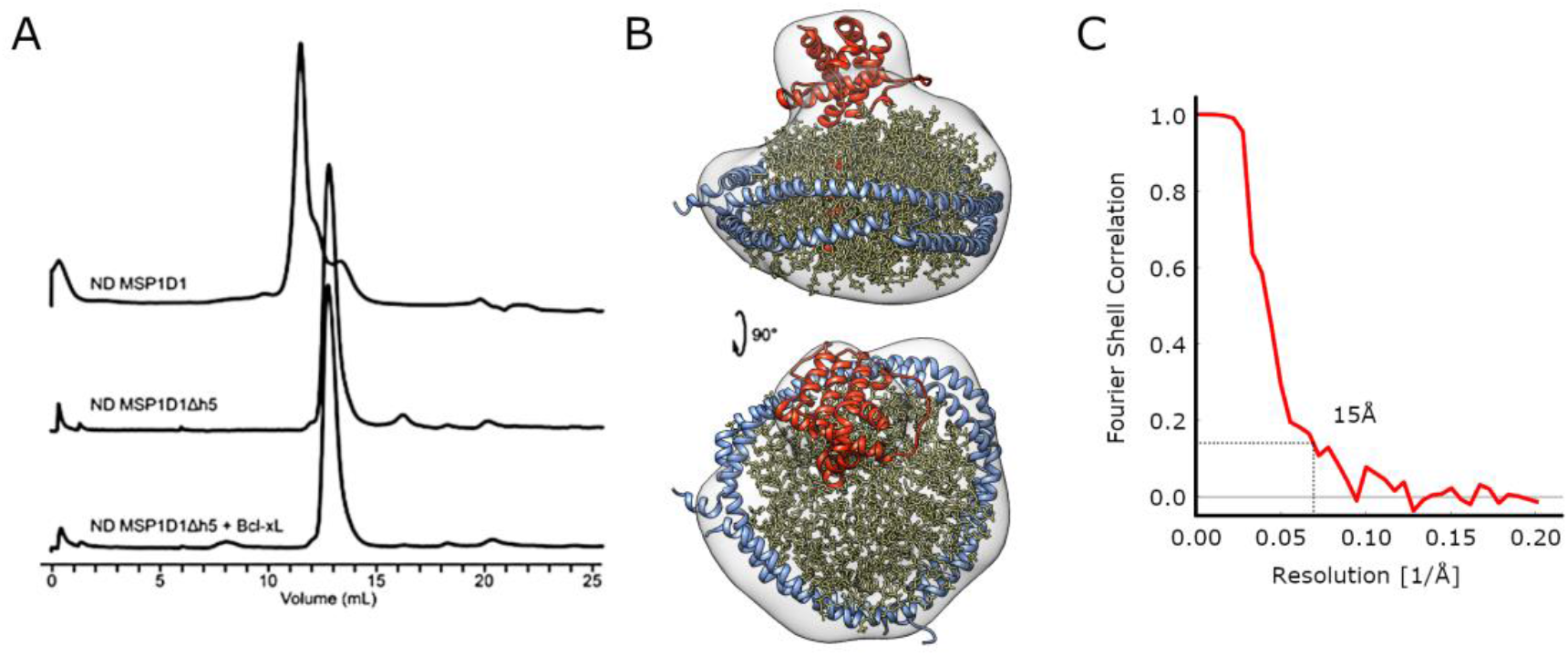
Structure of Bcl-xL embedded in MSP1D1Δh5 nanodiscs. (A) Size exclusion chromatography profiles of nanodiscs and nanodisc Bcl-xL assembly samples (B) Reconstruction with the model of Bcl-xL (red) and nanodiscs (lipids in yellow, protein belt in blue) fitted into the density. (C) Fourier Shell Correlation graph. The 0.143 cutoff indicates a resolution of 15 Å (dotted lines).

## CONCLUSIONS

We showed that the size distribution of nanodiscs can be derived directly from the data via the *F*_*max*_ signature of the sample and gives reliable indications for the homogeneity of the preparation. The analysis can essentially be run in real time and thus can be used to steer experimental data collection strategies. While the method described here targets disc-like objects, the Feret signature part of the analysis is agnostic to the underlying shape of the projected particles and can also be used to characterize the size distributions of arbitrary objects. A more complex Feret signature combining distributions of several Feret properties such as *F*_*min*_, *F*_*max*_ and Feret diameters obtained at 90° to the directions of the *F*_*min*_ *and F*_*max*_ can be used to characterize more complex shapes. One application would be the comparison of the experimental Feret signature with the Feret signature calculated from the reconstruction determined from a particular particle set to validate the structure or to detect contaminations in the particle set.

## ABREVIATIONS

*MSP*: membrane scaffold protein
*SMALPS*: steryne-maleic acid lipidic particles
*cND*: covalently circularized nanodiscs
*DLS*: dynamic light scattering
*F_max_*: maximum Feret diameter
*d_nd_*: estimated nanodisc diameter

## ACKNOWLEDGEMENTS

This work was supported by National Institutes of Health grant R01 5R01CA179087 (FMM, DH, NV), and the Fondation pour l’audition grant FPA RD-2019-14 (DH, NV). The authors thank Bertrand Raynal and Sébastien Brûlé from the Molecular Biophysics Facility at Institut Pasteur for help with biophysical analysis and Nicolas Wolff and his team for access to the akta purifier. The authors acknowledge the use of the Titan Krios, Tecnai Spirit T12 and auxiliary equipment at the cryo-EM unit of the Sanford Burnham Prebys Medical Discovery Institute, which was created in part with the support of US National Institutes of Health Grant S10-OD012372 (DH) and Pew Charitable Trust 864K625 innovation award funds (DH). The authors acknowledge the NanoImaging Core at Institut Pasteur for access to the Glacios and VitroBot. The NanoImaging Core was created with the help of a grant from the French Government’s Investissements d’Avenir program (EQUIPEX CACSICE - Centre d’analyse de systèmes complexes dans les environements complexes, ANR-11-EQPX-0008). The authors declare no competing financial interests.

